# Loss of CNOT9 begets impairment in gastrulation leading to embryonic lethality

**DOI:** 10.1101/2020.04.27.063172

**Authors:** Hemanta Sarmah, Kentaro Ito, Mari Kaneko, Takaya Abe, Tadashi Yamamoto

## Abstract

The multi-subunit eukaryotic CCR4-NOT complex imparts gene expression control primarily via messenger RNA (mRNA) decay. Here, we present the role of subunit CNOT9 in target mRNA decay during embryonic development. CNOT9 null mice appear normal by the onset of gastrulation (E7.0), however, exhibit growth and differentiation defects accompanied by extensive cell death by embryonic day 9.5 (E9.5). Sox-2 Cre conditional CNOT9 knockout mice show almost identical phenotype with brief delay in onset and progression, suggesting defects to be epiblast-dominant. Among various identified targets, we show that *Lefty2* mRNA expression is post-transcriptionally regulated by CNOT9. *Lefty2 3’-UTR* containing mRNA has significantly higher stability in cells expressing mutant form of CNOT9, relative to cells expressing wild-type CNOT9. In addition, CNOT9 primarily localizes within the cytoplasm and bridges interactions between the CCR4-NOT complex and miRNA-RISC complex in gastrulating embryos.

## Introduction

Messenger RNA (mRNA) decay is an indisputable component of gene expression regulation. In higher eukaryotes such as humans and mice, a conserved multi-subunit protein complex, known as CCR4-NOT, interacts with RNA binding proteins and collectively performs mRNA decay (Chapat and Corbo, 2014, Shirai et al., 2014, Collart, 2016, Miller and Reese, 2012). The complex is composed of a central scaffold CNOT1, regulatory subunits CNOT2 and CNOT3, a catalytic core consisting of CNOT6 or CNOT6L and CNOT7 or CNOT8 proteins, and three additional subunits CNOT9, CNOT10, and CNOT11 (Shirai et al., 2014). Precise molecular function of subunits CNOT9-11 is yet to be understood. In this study, we elucidate the molecular and physiological function of murine CNOT9 with regard to mechanisms on mRNA decay during embryonic development.

Embryonic development is a complex, yet fine-tuned orchestration of gene expression regulation (Reik, 2007, Tomancak et al., 2007, Wang et al., 2005, Yi et al., 2010). Germ layer differentiation, also known as gastrulation, is an event that takes place during embryonic day 6.5 to 8.5 of post-implantation mouse embryos (Gilbert, 2010). Cell lineage fates are determined during this stage based on various extracellular signaling molecules and chemical gradients (Basson, 2012, Hammerschmidt and Wedlich, 2008, Wang et al., 2012). With the help of targeted gene disruption methods in mice, various genes have been identified as key determinants indispensable for embryonic gastrulation. While a majority of studies have shown transcription regulation as the underlying mechanism by which these genes influence gastrulation, the post-transcriptional component has not been adequately investigated. In this study, we present a scenario of post-transcriptional regulation of a gastrulation related gene Lefty2, whose expression is controlled by CCR4-NOT complex subunit CNOT9.

In-vitro, CNOT9 (also known as Rqcd1) was first shown to mediate retinoic-acid (RA) induced differentiation of F9 teratocarcinoma cells by positively influencing *c-jun* transcription (Hiroi et al., 2002). In the same study, increased expression of CNOT9, as a result of RA treatment, was found to rather inhibit branching morphogenesis of embryonic lung, via unknown mechanisms (Hiroi et al., 2002). In other studies, CNOT9 was shown to have physical interactions with non CCR4-NOT components – c-Myb and NIF-1 proteins, resulting in transcription inhibition and ligand-dependent activation of RARs and RXRs, respectively (Garapaty et al., 2008, Haas et al., 2004). Recently, CNOT9 has been confirmed to be a core component of the CCR4-NOT complex through crystal structure analysis and has been implicated in microRNA mediated RNA decay (Mathys et al., 2014, Chen et al., 2014). Therefore, with such heterogeneity in knowledgebase around CNOT9, we chose to investigate its physiological relevance and molecular function. We characterized the phenotype exhibited by CNOT9 knock-out (KO) embryos, and its role in specific mRNA decay during gastrulation.

## Results

### Loss of CNOT9 causes defects in embryonic gastrulation

We generated *Cnot9*^+/LacZ^ mice as described in Materials and Methods (Fig.S1b). Although *Cnot9*^+/LacZ^ mice appeared phenotypically normal, upon crossing of male and female *Cnot9*^+/LacZ^ mice no *Cnot9*^LacZ/LacZ^ (KO) mice were born. Furthermore, no KO embryos could be genotyped at embryonic day 14.5 (E14.5), suggesting early embryonic lethality. To identify the timing of embryonic lethality and onset of phenotypic defects in KO embryos, we investigated gastrulation stage embryos, 7 to 10 days post coitum. During early Bud/Headfold stages (E7.25 to E7.75) and early somite stages (E7.75 to E8.0), no obvious differences in KO embryo morphology were observed when compared with wildtype (WT) and heterozygous (HE) littermates (Fig.1a,1b). However, during intermediate and late somite stages (E8.0 to E8.5), CNOT9 KO embryos displayed onset of phenotypic defects in the form of growth retardation of embryo-proper (Fig.1c). On the following embryonic day, these KO embryos showed severe growth defects characterized by small size, perturbed developmental pace indicated by delay in embryo curling, and lack of blood vessels within visceral yolk-sac regions (Fig. 1d). Contrary to the previous stage, KO embryos exhibited higher cleaved PARP expression compared to WT littermates, suggesting cell death within these embryos (Fig. 1d). Another day into development, KO embryos continued to display severe defects in terms of embryo size and morphology and showed a sustained increase in cleaved PARP expression compared to WT littermate controls (Fig. 1e). Mouse embryonic stem cells derived from KO blastocysts were viable and did not show any defects in terms of colony size, growth rates or expression of stemness markers, suggesting a gastrulation specific defect within KO embryos (data not shown).

**Figure 1:**
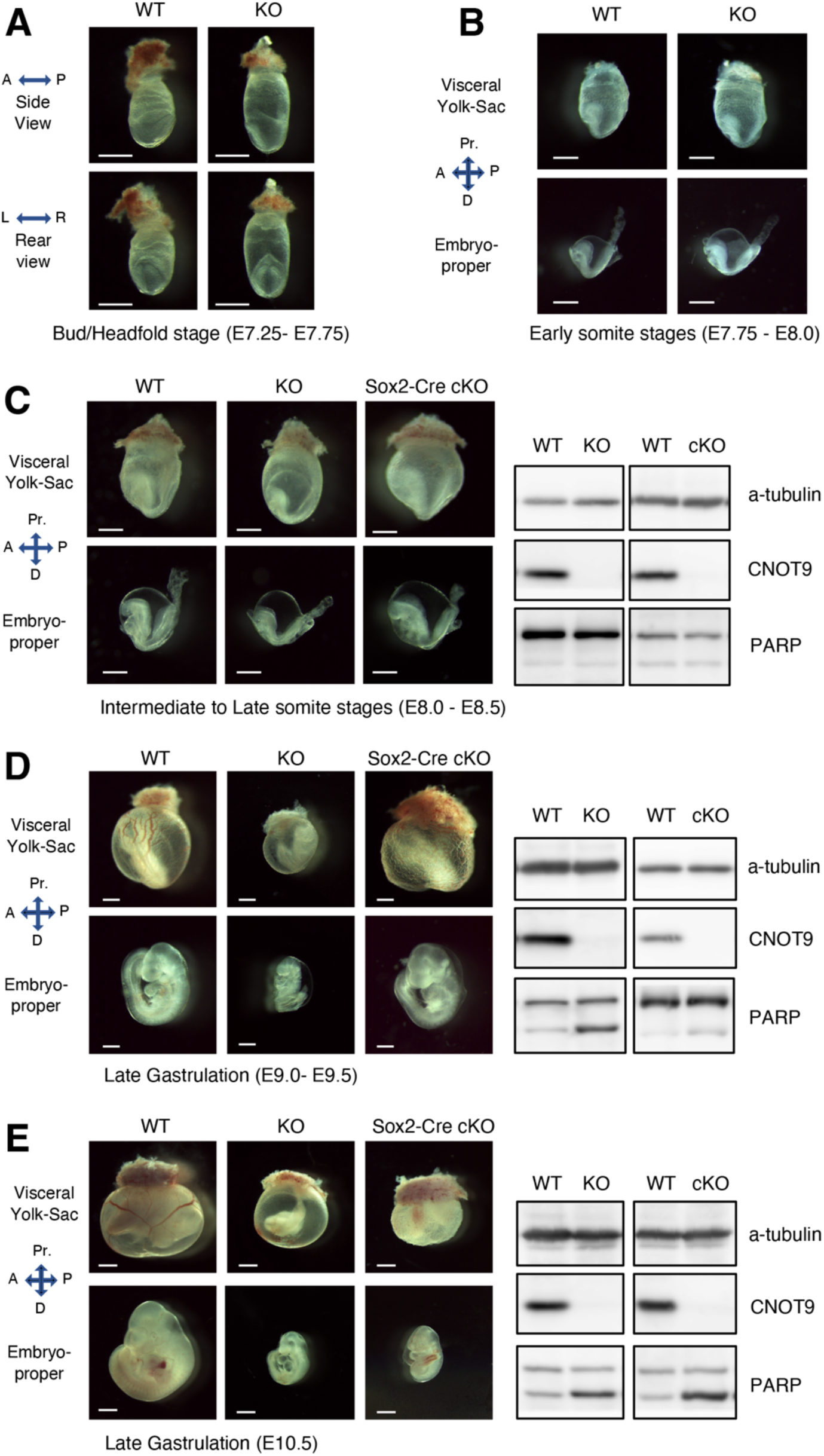
Embryo morphology and size evaluation of CNOT9 KO embryos alongside WT controls during (A) Bud and Headfold stages (E7.25-E7.75) (B) Early somite stages (E7.75 - E8.0) (C) Intermediate to Late Somite stages (E8.0 - E8.5) (D) and (E) Late gastrulation stages (E9.0 - E10.5). Western Blotting for PARP alongside corresponding stages indicates the extent of tissue death in KO embryos compared to WT control. Sox2-Cre cKO embryos indicate marginal delay in emergence of KO embryo phenotype. Scale: 500um

### CNOT9 KO embryo phenotype primarily contributed by epiblast lineage cells

Due to the high prevalence of placental defects in embryonic lethal mutant mice, we decided to test the contribution of trophoblast vs. epiblast lineage cells in driving KO embryo phenotype (Perez-Garcia et al., 2018). We performed histological analysis on placental regions during E8.5 and E9.5 stages and found that KO embryos possessed significantly smaller sized placentas compared to WT and HE littermate controls (Fig. S2a, S2b). A reduced or poorly developed placenta may, however, be caused due to an overall impairment in embryo development and differentiation. To test this observation from the standpoint of direct vs. indirect outcome of CNOT9 loss, we generated *Cnot9*^flox/flox^ mice, as described in Materials and Methods (Fig.S1a). Mice containing floxed *Cnot9* alleles were crossed with Sox2-Cre mice (B6.Cg-Edil3^Tg(Sox2-cre)1Amc^/J, Jackson Labs, USA) to generate epiblast specific knockout (eKO) mice. Similar to complete knockouts, no eKO mice were born suggesting embryonic lethality. Analysis of gastrulating eKO embryos during intermediate and late somite stages identified no characteristic morphological differences between WT controls (Fig. 1c). However, during late gastrulation stages, defects in eKO embryos began to surface, initially in regions of yolk-sac vasculature, followed by growth arrest and cell death in embryo-proper (Fig. 1c,1d).

### *Cnot9* expression during embryonic gastrulation via LacZ staining

Taking advantage of the LacZ knock-in allele in CNOT9 HE embryos, we performed b-galactosidase (b-gal) staining of embryos between E7.5 to E9.5 stage of gastrulation (Fig.2). At E7.5 stage, b-gal staining was predominantly seen within the epiblast region that would eventually contribute to form the embryo-proper (Fig.2a). At 8.5 stage, b-gal staining was seen in various parts of the embryo-proper that include the neuroepithelium, the trunk region, and the caudal region of the embryo. More importantly, b-gal staining was also seen in the extra-embryonic yolk-sac and within the ectoplacental cone (EPC) region of the embryo (Fig.2b). A day later, E9.5 stage embryos showed strong b-gal staining within labyrinthine and spongiotrophoblast regions of the placenta relative to weak and punctuated signal from regions of the embryo-proper (Fig.2c). A schematic representation of placental layers during E9.5 stage is depicted for visualization of lacZ expression within the placenta (Fig.2d). In addition to lacZ staining *Cnot9* mRNA expression (relative to *Gapdh*) at various regions of gastrulating embryos, was determined by qRT-PCR analysis (Fig.2e). In agreement with the b-gal staining pattern, *CNOT9* expression was about > 1.5 higher in E9.5 placenta compared to embryo-proper. Other stages showed nearly similar expression levels of *CNOT9* mRNA.

**Figure 2:**
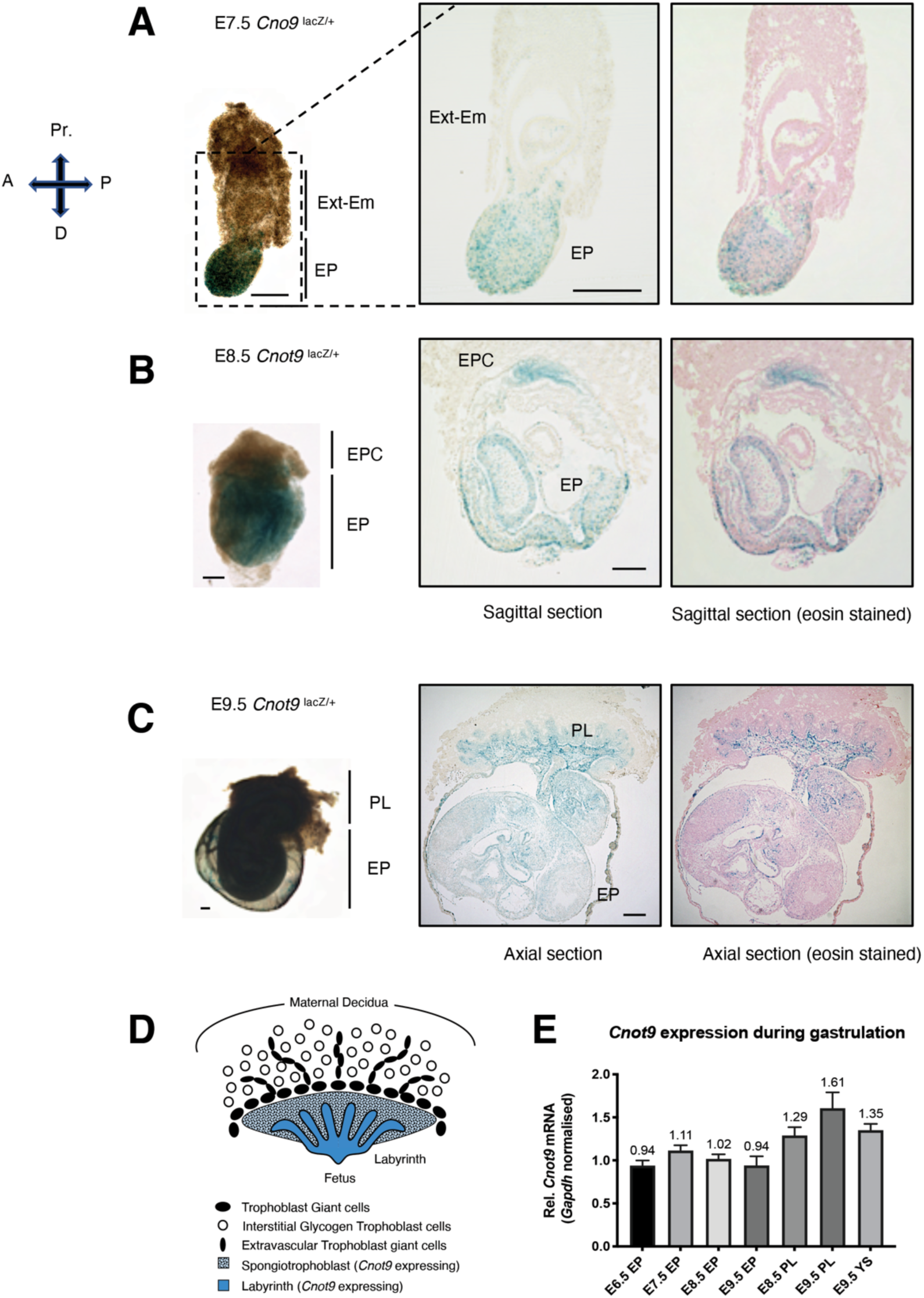
LacZ staining of CNOT9 HE embryos as a readout for spatial localization of *Cnot9* mRNA. **(A)** E7.5 stage indicates majority of expression in epiblast derived regions of the embryo. **(B)** E8.5 stage depicts expression in various regions of the embryo-proper with marginal expression in regions of the ectoplacental cone. **(C)** E9.5 staged embryos show extensive expression in regions of the placenta that correspond to labyrinth and spongiotrophoblast cells. Embryo-proper also shows expression although not uniform across the embryo. **(D)** Schematic representation depicting *Cnot9* expressing (LacZ stained) regions of the placenta in blue **(E)** *Cnot9* mRNA levels, relative to Gapdh, detected by qRT-PCR in WT embryos in various embryo regions across stages of gastrulation. Schematic representation Scale: 250um, Ext-Em: extra embryonic, EPC: Ectoplacental cone, EP: Embryo-proper, PL: Placenta, YS: Yolk-sac, A: Anterior end, P: Posterior end, Pr.: Proximal end, D: Distal end.

### Gene expression regulation via CNOT9

To investigate the molecular function of CNOT9 with added impetus from the KO embryo phenotype, we performed RNA-Seq analysis on three independent E8.0-E8.5 staged WT and KO samples. From a total of 15534 protein-coding targets, only 384 were upregulated >2-fold, while another 338 were downregulated >2-fold in KO embryos based on normalized RPKM scores (Fig.3a). Top few upregulated targets included *Lefty2, Lefty1, Nodal, Crypto-1*, and *Noto* that have previously been known for their role in embryonic gastrulation. A heatmap of a few select upregulated targets and downregulated targets are shown in Fig. 3b and Fig. S3a respectively. These in-silico results were further validated by qRT-PCR based analysis using E8.5 stage WT and KO cDNA samples (Fig. 3c). With the exception of *Noto*, other validated targets that showed upregulation at E8.5 stage, also showed either a trend or significant increase during E7.5 stage. This suggests that despite the lack of phenotypic defects during E7.5 stage, molecular events leading to abnormal embryo phenotype had already perpetuated within KO embryos. Primers used for the same have been listed in Fig. S3c. Pathway analysis performed using Gene-Set Enrichment Analysis (GSEA) algorithm clearly identified Nodal and TGF-beta signaling pathways to be affected in KO embryos, thereby summarizing the nature of gastrulation defects (Fig. 3d). Downregulated targets mainly suggested retarded embryo growth and differentiation characterized by significantly reduced expression levels of *Myl2, Epo, Fgf8, Sox10* and Pax family members - *Pax1, Pax5* and *Pax7* (Fig. S3b). Downregulated targets didn’t clearly suggest any specific signaling pathway (data not shown). In addition to upregulated targets involved in gastrulation, E8.5 stage KO embryos also showed higher expression for *Oct4* and *c-myb* mRNAs (Fig. 3c). We speculate that such an increase is contributed by a higher abundance of cells in an undifferentiated state within KO embryos, compared to controls.

**Figure 3:**
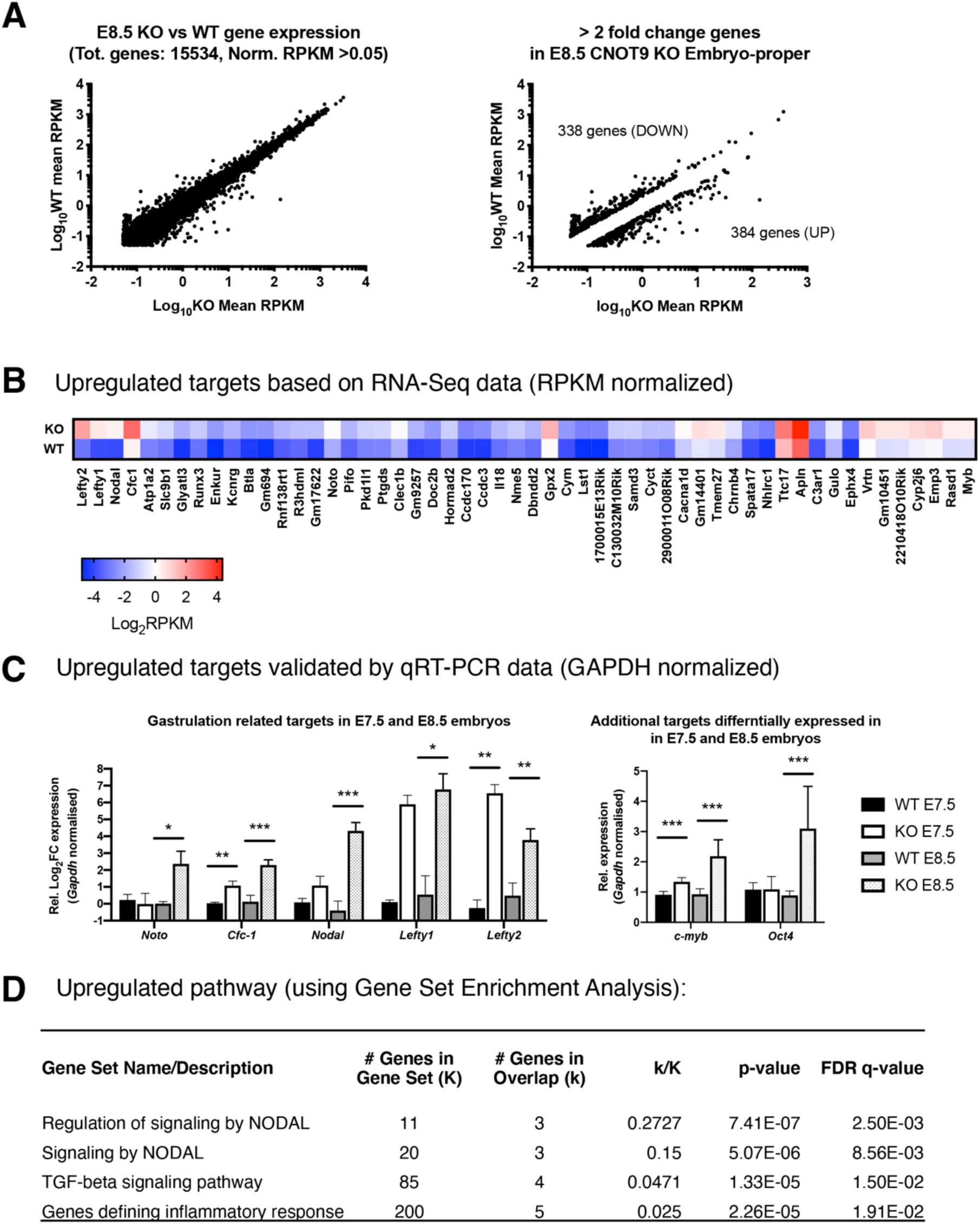
(A) Dot-plots representing gene expression in WT and KO embryos (left). More than 2-fold upregulated and downregulated targets (right) **(B)** Heat-map indicating average extent of upregulation for target genes in KO embryos compared to WT controls. **(C)** qRT-PCR based validation of upregulated genes involved in embryonic gastrulation in E7.5 and E8.5 stages (n=5, p < 0.05 (*), p < 0.01 (**), p < 0,001 (***). **(D)** Key upregulated pathways in CNOT9 KO embryos mainly involve TGF-beta signaling pathways influenced by Nodal and other competing ligands

### CNOT9 contributes to Lefty1/2 mRNA decay, in-vitro

In agreement with the current understanding of CNOT9 being associated with the mRNA decay complex, we found the protein to be predominantly localized within cytoplasmic fractions of gastrulating embryos (Fig. S4a). Furthermore, CNOT9 participated in complex formation with CCR4-NOT complex subunits - CNOT1, CNOT2, and CNOT3, as well as RISC component subunit GW182 during gastrulation as observed in a co-immunoprecipitation assay performed with anti-CNOT3 antibody. (Fig. S4b). To investigate the role of CNOT9 in the context of mRNA decay, we pursued two candidate targets - *Lefty1* and *Lefty2*, as validated in the previous section. Another important reason for picking these targets among others is the presence of miRNA binding sites within 3’UTR regions that is likely to facilitate mRNA decay, in-cis. Furthermore, we generated CNOT9 KO HeLa cells using CRISPR-Cas9 method for performing in-vitro biochemical assays. KO HeLa cells transfected with flag-hCNOT9 or flag-hCNOT9^mut4^ were used to test the decay kinetics of reporter luciferase conjugated with *Lefty2* or *Lefty1* 3’UTRs. ActinomycinD chase assay showed stabilization of both *Lefty2* or *Lefty1* 3’UTR conjugated reporter mRNAs in KO cells reconstituted with hCNOT9^mut4^ over a period of 6 hours (Fig 4a). Western blotting performed using anti-CNOT9 and anti-flag antibodies confirmed equal dose of flag tagged CNOT9 protein expression (Fig 4c). *c-myb* 3’UTR known to possess multiple microRNA binding sites, clearly showed impaired reporter mRNA decay in the absence of CNOT9 (Fig 4a). *p21(cdkn1a)* 3’UTR conjugated reporter, used as a negative control, did not show any significant difference in decay kinetics (Fig 4b). Similar decay kinetics for hFHL3 mRNA (endogenous control) showed uniformity of ActinomycinD treatment across different samples, over the course of the experiment.

**Figure 4:**
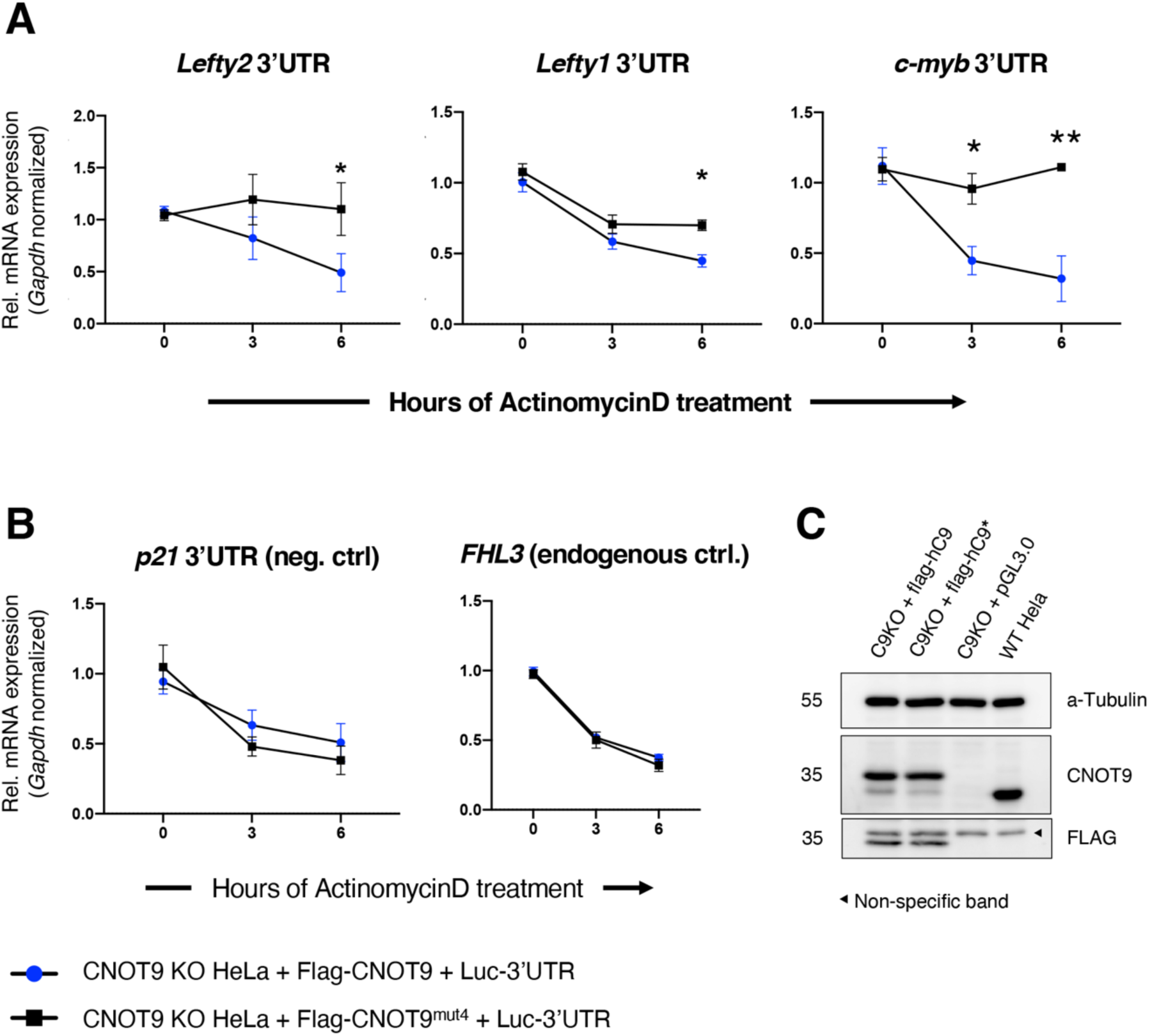
Decay kinetics of Luciferase reporter conjugated with 3’UTR elements of (A) *Lefty1, Lefty2*, and *c-myb* (B) *p21* 3’UTR as non-target control and *Fhl3* as endogenous control. Blue dots represent relative expression of reporter mRNA in CNOT9 KO cells reconstituted with human CNOT9, whereas black dots represent expression in CNOT9 KO cells reconstituted with mutant CNOT9 (CNOT9^mut4^) that cannot interact with CNOT1. (C) Western Blotting using anti-CNOT9 and anti-FLAG antibodies to ensure equal expression of CNOT9 and CNOT9^mut4^ proteins. Values are Mean ± SD [n=3, p < 0.05 (*), p < 0.01 (**)]

## Discussion

The mammalian CCR4-NOT complex is profoundly involved in embryonic development. For instance, loss of CNOT3 leads to pre-implantation growth arrest in mouse embryos (Neely et al., 2010, Morita et al., 2011). In this study we highlight the physiological role of CNOT9 during embryonic gastrulation. *Cnot9* knockout mice exhibited developmental abnormalities during mid-gastrulation stages (E8.0 to E8.5) leading to embryonic lethality at E9.5 stage. KO embryos showed a spectrum of defects that included reduced size, arrested growth, pale colorization of embryo-proper, impaired yolk-sac vasculature, and significantly reduced placenta. Chorioallantoic fusion defects were not observed in E8.5 KO embryos (data not shown). To determine the weightage contribution of epiblast vs. trophoblast derived cells, we examined the phenotype of embryos that underwent epiblast specific depletion of CNOT9. A marginal delay in the onset of phenotype and timing of embryo lethality was observed in Sox2-Cre conditional knockout mice compared to complete knockouts. This suggested that phenotypic defects in KO embryos were mainly contributed by cells originating from the epiblast lineage. In other words, a wildtype trophoblast lineage could not profoundly rescue the phenotype observed in whole-body knockouts. As a first step towards examining function, we investigated the expression pattern of *Cnot9* mRNA using whole-mount LacZ staining and qRT-PCR based analysis. Initially (E7.5), *Cnot9* was almost exclusively expressed within the epiblast, but late in gastrulation, (E8.5 to E9.5) expression was also seen within placental regions. It is worth noting that both E8.5 and E9.5 placental regions showed higher expression of *Cnot9* compared to relative levels within the adjoining embryo proper. Unfortunately, due to the timing and nature of defects, both KO and conditional knockout mice models limit our understanding of CNOT9 within a functional placenta. It would be worth investigating CNOT9 function within placental lineage via in-vitro trophoblast stem cell-based differentiation assays, or in-vivo models that produce placenta specific depletion.

Phenotypic defects in KO embryos motivated us to address the molecular function of CNOT9 in developmental context. E8-E8.5 staged WT and KO embryos were pooled and subjected to RNA-Seq analysis. With biochemical validation, we observed significant upregulation in mRNA levels of *Lefty1/2, Nodal, Cfc-1*, among others that have previously been shown to have profound roles in embryonic development (Meno et al., 1998, Meno et al., 1999, Conlon et al., 1994, Yan et al., 1999). For instance, overexpression of Lefty2 in zebrafish embryos blocks head and trunk mesoderm formation (Meno et al., 1999). *Cfc-1* overexpression in mice caused downregulation of genes involved in cardiac differentiation during mid-gastrulation stages whereas, in chick embryos, its gain-of-function led to suppression of posterior mesodermal fates while promoting anterior mesoderm development (Chu et al., 2005, Lin et al., 2016). Nodal overexpression has been shown to promote differentiation of mouse ES cells into endoderm and mesoderm lineages at the expense of neuroectoderm formation (Pfendler et al., 2005). Despite these findings, little is known from the standpoint of post-transcriptional regulation of such transcripts and their consequences on embryo development. One such report showed *Lefty2* mRNA regulation via microRNA-127 in mouse ES cells (Ma et al., 2016). Therefore, we present a model where the physiological relevance of the upregulation of gastrulation related transcripts can be studied.

We focused on *Lefty1* and *Lefty2* transcripts mainly because of the extent of upregulation and the presence of canonical microRNA binding sites in their 3’UTR elements. With the help of TargetScan database, we found that both mRNAs contain an AAGCACU element that serves as binding sites for miR-291-3p, miR-294-3p, miR-295-3p, miR-302-3p microRNA. Therefore, we cloned 3’UTR elements in pGL3.0 luciferase expression vectors. In WT HeLa cells, relative to vector-only control, luciferase reporter expression and activity were significantly reduced when conjugated with *Lefty1* and *Lefty2* 3’UTRs (data not shown). We further clarified that in absence of wild-type human CNOT9, mRNA decay of reporters containing *Lefty1* or *Lefty2* 3’UTRs was significantly affected. Due to very high sequence conservation of human vs. mouse Cnot9 proteins (99.7%), and microRNA binding sites within UTR elements of mouse and human *Lefty* transcripts, a similar mechanism is likely to be present in humans.

The pale color of E8.5 - E9.5 stage KO embryos can be explained in terms of dysfunctional waves of primitive and definitive waves of erythropoiesis depending on *c-myb* expression. Downregulation of *c-myb* transcription has previously been reported to be imperative for erythropoietin (epo) induced differentiation (Todokoro et al., 1988). Furthermore, retinoic acid has been shown to suppress *c-myb* expression (Mandelbaum et al., 2018). CNOT9 has previously been proposed to be retinoic acid-inducible and therefore, in agreement with these ideas, our data suggests a strong correlation in terms of loss of CNOT9, the elevation of *c-myb* expression, and reduced *Epo* expression (Hiroi et al., 2002). The difference, however, lies in the manner in which c-myb mRNA expression is elevated. These two evidences showed transcriptional control in *c-myb* expression, while our data suggests post-transcriptional control (via mRNA stabilization) in the backdrop of CNOT9 loss. Although defects in blood formation may be triggered by improper germ layer differentiation at earlier stages of gastrulation, we speculate that elevation of *c-myb* mRNA can inhibit *epo* induced erythroid differentiation in CNOT9 KO or conditional knockout mice. Further analysis using mice models for erythroid-specific deletion can help elucidate this aspect of CNOT9 function.

CNOT9 has been proposed to play regulatory roles within the complex in a manner that assists mRNA decay by interacting with CNOT1 and stimulating catalysis by subunits CNOT6, CNOT6L, CNOT7 and CNOT8 (Pavanello et al., 2018).This idea was further perpetuated by an in-vitro study of the human CCR4-NOT complex, which suggested that stimulus-dependent CNOT9 interaction with RNA binding proteins results in targeted mRNA decay (Raisch et al., 2019). In congruity with this idea we observed a similar trend within our RNA-Seq data. Only 722 (less than 5%) out of 15534 total detected protein-coding genes underwent more than 2-fold significantly change. In other words, developmental stimulus affects the expression of very specific targets via CNOT9, leaving behind a majority of them unaffected.

The molecular function of CNOT9 in regulating target mRNA decay either by microRNA pathway or via interactions with RNA binding proteins, demands further investigation. Recently, two studies had shown that interactions between GW182 proteins with CNOT1-CNOT9 complex facilitate miRNA mediated decay (Chen et al., 2014, Mathys et al., 2014). Their findings suggested that the presence of CNOT9 provided structural stability to the trimeric complex and augmented the affinity of GW182 for CNOT1. Because their experiments were performed either in cell-free conditions or in wild-type HEK-293T cells that may express endogenous levels of CNOT9, we tested this theory using CNOT9 KO HeLa cells. We found that absence of CNOT9 completely abrogated interactions between the C-terminal silencing domain of GW182 with CCR4-NOT subunits CNOT1 and CNOT3 (Fig. S5). This makes CNOT9 an indispensable element within the molecular framework of microRNA mediated mRNA decay.

## Materials and Methods

### Generation of Cnot9-knockout and Cnot9 conditional mice

Targeting vector used for generation of *Cnot9*-knockout mice (Accession. No. CDB0573K: http://www2.clst.riken.jp/arg/mutant%20mice%20list.html) was constructed from a 129SVJ genomic clone (STRATAGENE, CA). Exon 1 of the *Cnot9* locus was targeted and replaced by *LacZ* and *neomycin* resistance gene by electroporating TT2 ES cells (Yagi et al., 1993). Neomycin-resistant ES cell clones harboring the homologous recombination were screened by PCR and Southern blot analysis. Two correctly targeted ES cell lines were injected into ICR 8-cell-stage embryos to generate chimeric mice. Germline-transmitted male chimeras were crossed with C57BL/6J female mice (Japan CLEA, Tokyo) to obtain heterozygous F1 offspring. Genotyping was performed using: 5’-TTATCTGGACGCGGGTTGTGAATGCTGG-3’ (Pr1), 5’-ACTAGTTCTAGAGCGGCCGATTTAAATACG-3’ (Pr2) and 5’-GTCCTAAGAAAGACATTCCAGGTAGAG-3’ (Pr3) (Fig.S1c). Cnot9 conditional KO (Cnot9^fl/fl^) mice (Accession. No. CDB1112K) were also generated similarly by targeting TT2 ES cell lines. To generate conditional alleles (floxed alleles) from targeted alleles, mice with targeted alleles were crossed with mice expressing FLP (Jackson #009086). Genotyping was performed using: 5’-CATGGGCTCATTAGCTGTCAAACAGGTTGAG-3’ (Pr1), and 5’-CCACTGATAGATCTTCTCTCTGTCCACTTGG-3’ (Pr2) (Fig.S1d). Experiments were performed with mice that had been backcrossed successively to C57BL/6J mice for at least seven generations. Animals were maintained in a 12-hour light dark-cycle within a temperature-controlled (22°C) barrier facility with abundant food (Rodent Diet CA-1, CLEA Japan) and water. All animal experiments were carried out following guidelines for animal use issued by Animal Resources Section, OIST Graduate University and the Institutional Animal Care and Use Committee (IACUC), RIKEN Kobe Branch.

### Antibodies

Antibodies against CNOT1 and CNOT3 have been previously described (Chen et al., 2011). Commercially available antibodies used in this study were: anti-a-tubulin (Sigma, T9026), anti-PARP (CST, 9542S), anti-CNOT9 (Proteintech, 22503-1-AP), anti-Flag (MBL, PM020), anti-Xrn-1(Bethyl A300-443A), anti-HistoneH3 (CST, 4499S), anti-GW182 (Bethyl, A302-329A), anti-CNOT2 (CST, 34214S), anti-4EBP1 (CST, 9644S), anti-b-actin (CST, 4970L), anti-GAPDH (CST, 2118L), anti-CNOT10 (Bethyl A304-899A), and anti-Ago2 (CST, 2897S).

### Tissue staining and histology

Post dissection, placental tissue was fixed in 4% paraformaldehyde (PFA) solution overnight, followed by stepwise dehydration in Ethanol solutions and finally embedded in paraffin. 8-10μm sections generated using Microm HM325 rotary microtome were stained with Hematoxylin 3G (8656) and Eosin solutions (8659) (Sakura Finetek, Japan). For LacZ staining, E7.5, E8.5, and E9.5 stage embryos were first fixed in ice-cold 0.2% Glutaraldehyde solution of a period of 5, 10 and 15 minutes respectively, followed by X-gal staining at 37°C for 48 hours. Whole-mount LacZ stained embryos were re-fixed in 4% PFA solution overnight, embedded in JB4 resin (Polysciences, Inc.) and sectioned (10um) using Microtome Rotatif, HM335E.

### Mouse embryo and tissue section imaging

All images for mouse embryos across various stages were taken using Leica IC80 HD stereomicroscope. Tissue sections of embryo and placenta were imaged using Keyence BZ-X710 microscope.

### Plasmids

Flag-tagged human CNOT9 was cloned into pCDNA3.0 and CNOT9^mut4^ (H58A F60A A64Y V71Y) was generated using point mutagenesis. Mouse Lefty1 3′-UTR (425 bp), Lefty2 3’-UTR (1290 bp), c-myb 3’-UTR (1243 bp), were cloned into the pGL3 control vector. p21 3’-UTR in pGL3.0 control vector was generated by Dr. Akinori Takahashi in the laboratory. Flag-tagged full-length silencing domain of human GW182/TNRC6C (1260-1690 amino acids) was PCR amplified from RIKEN cDNA library (KIAA1582) and cloned into pCDNA3.0 vector.

### Generation of CNOT9 KO Hela cells with CRISPR/Cas9

Guide RNA sequences were designed using CHOP-CHOP website, provided by the University of Bergen, Norway (Labun et al., 2019). Insert oligonucleotides targeting exon1 of hCNOT9 were 5’-CACCGGCACAGCCTGGCGACGGCTG-3’ and 5’-AAACCAGCCGTCGCCAGGCTGTGCC-3’. These complementary oligos were annealed and cloned into pSpCas9(BB)-2A-Puro vector (Addgene). Plasmids containing guideRNA target sites were confirmed by sequencing. HeLa cells were transfected using pSpCas9(BB)-2A-Puro vectors containing guide RNAs that target hCNOT9 (test) or EGFP (control), as described previously (Ran et al., 2013). 24 hours post transfection, cells were treated with 2.5 μg/ml of puromycin for three days. After 3 days, cells were collected, and seeded in three 10 cm plates at a density of 300 cells per dish. After 10-12 days of culture, small colonies were picked and seeded into individual wells of a 48 well plate and propagated further. Knockout clones were confirmed by DNA sequencing and Western blotting.

### Cell Culture

HeLa cells were cultured in DMEM (low Glucose) containing 10% FBS. TransIT-LT1 (Mirus Bio) reagent was used for transient transfection assays based on manufacturer’s protocol. For determination of reporter mRNA half-life kinetics CNOT9 KO HeLa cells were first transfected with flag-hCNOT9 or flag-hCNOT9^mut4^, for 24 hours, followed by transfection of luciferase-*Lefty2* 3′-UTR mRNA, luciferase-*Lefty1* 3′-UTR mRNA, or luciferase-*c-myb* 3′-UTR mRNA for 4 hours. Cells were then treated with 2.5ug/ml ActinomycinD (Wako) for 6 hours with samples being collected at 0, 3 and 6 hours after treatment.

### Quantitative RT-PCR

Total RNA was isolated from mouse embryos and HeLa cells using Isogen II reagent (Nippongene), followed by cDNA synthesis using with SuperScript Reverse Transcriptase III Kit (Invitrogen), based on manufacturers protocol. For real-time PCR, cDNA (1:10 diluted) was mixed with target-specific primers and SYBR Green Supermix (Takara). Data analysis was done using Viia7 sequence detection interface (Applied Biosystems). mRNA expression for various targets was determined relative to mouse or human GAPDH levels using ΔΔCt method.

### Western Blotting

Western Blotting for various proteins/targets was done via detection of chemiluminescence (Amersham Bioscience) as described previously (Takahashi et al., 2015). Embryos or cultured cells were solubilized in TNE buffer (50 mM Tris-HCl [pH 7.5], 150 mM NaCl, 1 mM EDTA, 1% NP40, and 1 mM PMSF) for 10 minutes at 4°C, after performing 10-second pulse homogenization of pellet (Fisherbrand Pellet Pestle). Total protein concentration was determined using BCA assay (Thermo Fisher), and adjusted by addition of appropriate amount of SDS sample buffer to lysates. Lysates were subjected to SDS-PAGE electrophoresis and electro-transferred into PVDF (polyvinylidene difluoride) membranes. Bands corresponding to various proteins were detected by corresponding antibodies using ImageQuant LAS 4000 (GE Healthcare, Tokyo) and analyzed using ImageQuant software. Nuclear and cytoplasmic fractions were isolated using the NE-PER kit (Thermo Scientific 78833) according to the manufacturer’s protocol.

### Protein Immunoprecipitation

E8.5 embryo lysates were subjected to immunoprecipitation using anti-CNOT3 antibody or mouse control IgG (Santacruz, sc-2025) for 1 hour at 4°C, followed by an additional 1-hour incubation with 50μl of Dynabeads (Invitrogen) added to the antibody-lysate solution. Immunoprecipitated proteins were recovered by magnetic separation of bead-antibody-protein complex. Subsequently, individual proteins in immune complexes were immunoblotted with appropriate antibodies. HeLa cell lysates obtained after transfecting cells with flag-hCNOT9 or flag-hCNOT9^mut4^ or pCDNA3.0 vectors were subjected to immunoprecipitation using ANTI-FLAG M2 Affinity gel (Sigma) for 2 hours at 4°C. All lysates were subjected to BCA assay prior to IP, to ensure equal amount of total protein.

### RNA-Seq analysis

High-quality total RNA (500ng) was isolated from pooled E8.0-E8.5 stage embryos using Isogen II reagent (Nippongene) and subjected to RNA-seq library preparation using TruSeq Stranded mRNA LT Sample Prep Kit (Illumina) based on manufacturer’s protocol. Paired-end RNA sequencing (109 base-pair) with a Hiseq PE Rapid Cluster Kit v2-HS and Hiseq Rapid SBS Kit v2-HS (200 Cycle) on a Hiseq2500 (Illumina) was performed following manufacturer’s protocol. Sequencing reads were mapped to mouse reference genome provided in Ensembl database using StrandNGS software (Strand Life Sciences). Data analysis was done by mapping reads to the mm10 genome sequence (Ensembl), thereby converting raw counts to corresponding FPKM scores. For downstream analysis, only protein-coding genes with FPKM scores ≥ 0.05 were selected. Gene set enrichment analysis (GSEA) was performed using online software from Broad Institute, USA. Sequence data are available through ArrayExpress under the accession number (****).

### Statistical analysis

Unpaired, two-tailed Student’s t-test was used for data analysis. Bar graphs represent mean ± standard error of the means (SEM) and p-value < 0.05 was deemed statistically significant.

## Notes

### Competing Interest Statement

The authors have declared no competing interest.

## References

Basson, M. A. 2012. Signaling in cell differentiation and morphogenesis. Cold Spring Harb Perspect Biol, 4.

Chapat, C. & Corbo, L. 2014. Novel roles of the CCR4-NOT complex. Wiley Interdiscip Rev RNA, 5, 883–901.

Chen, Y., Boland, A., Kuzuoglu-Ozturk, D., Bawankar, P., Loh, B., Chang, C. T., Weichenrieder, O. & Izaurralde, E. 2014. A DDX6-CNOT1 complex and W-binding pockets in CNOT9 reveal direct links between miRNA target recognition and silencing. Mol Cell, 54, 737–50.

Chu, J., Ding, J., Jeays-Ward, K., Price, S. M., Placzek, M. & Shen, M. M. 2005. Non-cell-autonomous role for Cripto in axial midline formation during vertebrate embryogenesis. Development, 132, 5539–51.

Collart, M. A. 2016. The Ccr4-Not complex is a key regulator of eukaryotic gene expression. Wiley Interdiscip Rev RNA, 7, 438–54.

Conlon, F. L., Lyons, K. M., Takaesu, N., Barth, K. S., Kispert, A., Herrmann, B. & Robertson, E. J. 1994. A primary requirement for nodal in the formation and maintenance of the primitive streak in the mouse. Development, 120, 1919–28.

Garapaty, S., Mahajan, M. A. & Samuels, H. H. 2008. Components of the CCR4-NOT complex function as nuclear hormone receptor coactivators via association with the NRC-interacting Factor NIF-1. J Biol Chem, 283, 6806–16.

Gilbert, S. F. 2010. Developmental biology, Sunderland, Sinauer.

Haas, M., Siegert, M., Schurmann, A., Sodeik, B. & Wolfes, H. 2004. c-Myb protein interacts with Rcd-1, a component of the CCR4 transcription mediator complex. Biochemistry, 43, 8152–9.

Hammerschmidt, M. & Wedlich, D. 2008. Regulated adhesion as a driving force of gastrulation movements. Development, 135, 3625–41.

Hiroi, N., Ito, T., Yamamoto, H., Ochiya, T., Jinno, S. & Okayama, H. 2002. Mammalian Rcd1 is a novel transcriptional cofactor that mediates retinoic acid-induced cell differentiation. EMBO J, 21, 5235–44.

Labun, K., Montague, T. G., Krause, M., Torres Cleuren, Y. N., Tjeldnes, H. & Valen, E. 2019. CHOPCHOP v3: expanding the CRISPR web toolbox beyond genome editing. Nucleic Acids Res, 47, W171–W174.

Lin, X., Zhao, W., Jia, J., Lin, T., Xiao, G., Wang, S., Lin, X., Liu, Y., Chen, L., Qin, Y., Li, J., Zhang, T., Hao, W., Chen, B., Xie, R., Cheng, Y., Xu, K., Yao, K., Huang, W., Xiao, D. & Sun, Y. 2016. Ectopic expression of Cripto-1 in transgenic mouse embryos causes hemorrhages, fatal cardiac defects and embryonic lethality. Sci Rep, 6, 34501.

Ma, H., Lin, Y., Zhao, Z. A., Lu, X., Yu, Y., Zhang, X., Wang, Q. & Li, L. 2016. MicroRNA-127 Promotes Mesendoderm Differentiation of Mouse Embryonic Stem Cells by Targeting Left-Right Determination Factor 2. J Biol Chem, 291, 12126–35.

Mandelbaum, J., Shestopalov, I. A., Henderson, R. E., Chau, N. G., Knoechel, B., Wick, M. J. & Zon, L. I. 2018. Zebrafish blastomere screen identifies retinoic acid suppression of MYB in adenoid cystic carcinoma. J Exp Med, 215, 2673–2685.

Mathys, H., Basquin, J., Ozgur, S., Czarnocki-Cieciura, M., Bonneau, F., Aartse, A., Dziembowski, A., Nowotny, M., Conti, E. & Filipowicz, W. 2014. Structural and biochemical insights to the role of the CCR4-NOT complex and DDX6 ATPase in microRNA repression. Mol Cell, 54, 751–65.

Meno, C., Gritsman, K., Ohishi, S., Ohfuji, Y., Heckscher, E., Mochida, K., Shimono, A., Kondoh, H., Talbot, W. S., Robertson, E. J., Schier, A. F. & Hamada, H. 1999. Mouse Lefty2 and zebrafish antivin are feedback inhibitors of nodal signaling during vertebrate gastrulation. Mol Cell, 4, 287–98.

Meno, C., Shimono, A., Saijoh, Y., Yashiro, K., Mochida, K., Ohishi, S., Noji, S., Kondoh, H. & Hamada, H. 1998. lefty-1 is required for left-right determination as a regulator of lefty-2 and nodal. Cell, 94, 287–97.

Miller, J. E. & Reese, J. C. 2012. Ccr4-Not complex: the control freak of eukaryotic cells. Crit Rev Biochem Mol Biol, 47, 315–33.

Morita, M., Oike, Y., Nagashima, T., Kadomatsu, T., Tabata, M., Suzuki, T., Nakamura, T., Yoshida, N., Okada, M. & Yamamoto, T. 2011. Obesity resistance and increased hepatic expression of catabolism-related mRNAs in Cnot3+/-mice. EMBO J, 30, 4678–91.

Neely, G. G., Kuba, K., Cammarato, A., Isobe, K., Amann, S., Zhang, L., Murata, M., Elmen, L., Gupta, V., Arora, S., Sarangi, R., Dan, D., Fujisawa, S., Usami, T., Xia, C. P., Keene, A. C., Alayari, N. N., Yamakawa, H., Elling, U., Berger, C., Novatchkova, M., Koglgruber, R., Fukuda, K., Nishina, H., Isobe, M., Pospisilik, J. A., Imai, Y., Pfeufer, A., Hicks, A. A., Pramstaller, P. P., Subramaniam, S., Kimura, A., Ocorr, K., Bodmer, R. & Penninger, J. M. 2010. A global in vivo Drosophila RNAi screen identifies NOT3 as a conserved regulator of heart function. Cell, 141, 142–53.

Pavanello, L., Hall, B., Airhihen, B. & Winkler, G. S. 2018. The central region of CNOT1 and CNOT9 stimulates deadenylation by the Ccr4-Not nuclease module. Biochem J, 475, 3437–3450.

Perez-Garcia, V., Fineberg, E., Wilson, R., Murray, A., Mazzeo, C. I., Tudor, C., Sienerth, A., White, J. K., Tuck, E., Ryder, E. J., Gleeson, D., Siragher, E., Wardle-Jones, H., Staudt, N., Wali, N., Collins, J., Geyer, S., Busch-Nentwich, E. M., Galli, A., Smith, J. C., Robertson, E., Adams, D. J., Weninger, W. J., Mohun, T. & Hemberger, M. 2018. Placentation defects are highly prevalent in embryonic lethal mouse mutants. Nature, 555, 463–468.

Pfendler, K. C., Catuar, C. S., Meneses, J. J. & Pedersen, R. A. 2005. Overexpression of Nodal promotes differentiation of mouse embryonic stem cells into mesoderm and endoderm at the expense of neuroectoderm formation. Stem Cells Dev, 14, 162–72.

Raisch, T., Chang, C. T., Levdansky, Y., Muthukumar, S., Raunser, S. & Valkov, E. 2019. Reconstitution of recombinant human CCR4-NOT reveals molecular insights into regulated deadenylation. Nat Commun, 10, 3173.

Ran, F. A., Hsu, P. D., Wright, J., Agarwala, V., Scott, D. A. & Zhang, F. 2013. Genome engineering using the CRISPR-Cas9 system. Nat Protoc, 8, 2281–2308.

Reik, W. 2007. Stability and flexibility of epigenetic gene regulation in mammalian development. Nature, 447, 425–32.

Shirai, Y. T., Suzuki, T., Morita, M., Takahashi, A. & Yamamoto, T. 2014. Multifunctional roles of the mammalian CCR4-NOT complex in physiological phenomena. Front Genet, 5, 286.

Takahashi, A., Adachi, S., Morita, M., Tokumasu, M., Natsume, T., Suzuki, T. & Yamamoto, T. 2015. Post-transcriptional Stabilization of Ucp1 mRNA Protects Mice from Diet-Induced Obesity. Cell Rep, 13, 2756–67.

Todokoro, K., Watson, R. J., Higo, H., Amanuma, H., Kuramochi, S., Yanagisawa, H. & Ikawa, Y. 1988. Down-regulation of c-myb gene expression is a prerequisite for erythropoietin-induced erythroid differentiation. Proc Natl Acad Sci U S A, 85, 8900–4.

Tomancak, P., Berman, B. P., Beaton, A., Weiszmann, R., Kwan, E., Hartenstein, V., Celniker, S. E. & Rubin, G. M. 2007. Global analysis of patterns of gene expression during Drosophila embryogenesis. Genome Biol, 8, R145.

Wang, J., Sinha, T. & Wynshaw-Boris, A. 2012. Wnt signaling in mammalian development: lessons from mouse genetics. Cold Spring Harb Perspect Biol, 4.

Wang, S., Cowan, C. A., Chipperfield, H. & Powers, R. D. 2005. Gene expression in the preimplantation embryo: in-vitro developmental changes. Reprod Biomed Online, 10, 607–16.

Yagi, T., Tokunaga, T., Furuta, Y., Nada, S., Yoshida, M., Tsukada, T., Saga, Y., Takeda, N., Ikawa, Y. & Aizawa, S. 1993. A novel ES cell line, TT2, with high germline-differentiating potency. Analytical biochemistry, 214, 70–6.

Yan, Y. T., Gritsman, K., Ding, J., Burdine, R. D., Corrales, J. D., Price, S. M., Talbot, W. S., Schier, A. F. & Shen, M. M. 1999. Conserved requirement for EGF-CFC genes in vertebrate left-right axis formation. Genes Dev, 13, 2527–37.

Yi, H., Xue, L., Guo, M. X., Ma, J., Zeng, Y., Wang, W., Cai, J. Y., Hu, H. M., Shu, H. B., Shi, Y. B. & Li, W. X. 2010. Gene expression atlas for human embryogenesis. FASEB J, 24, 3341–50.

