## Supplementary Figures and Tables for "Loss of CNOT9 begets impairment in gastrulation leading to embryonic lethality"

### Supplementary Information:

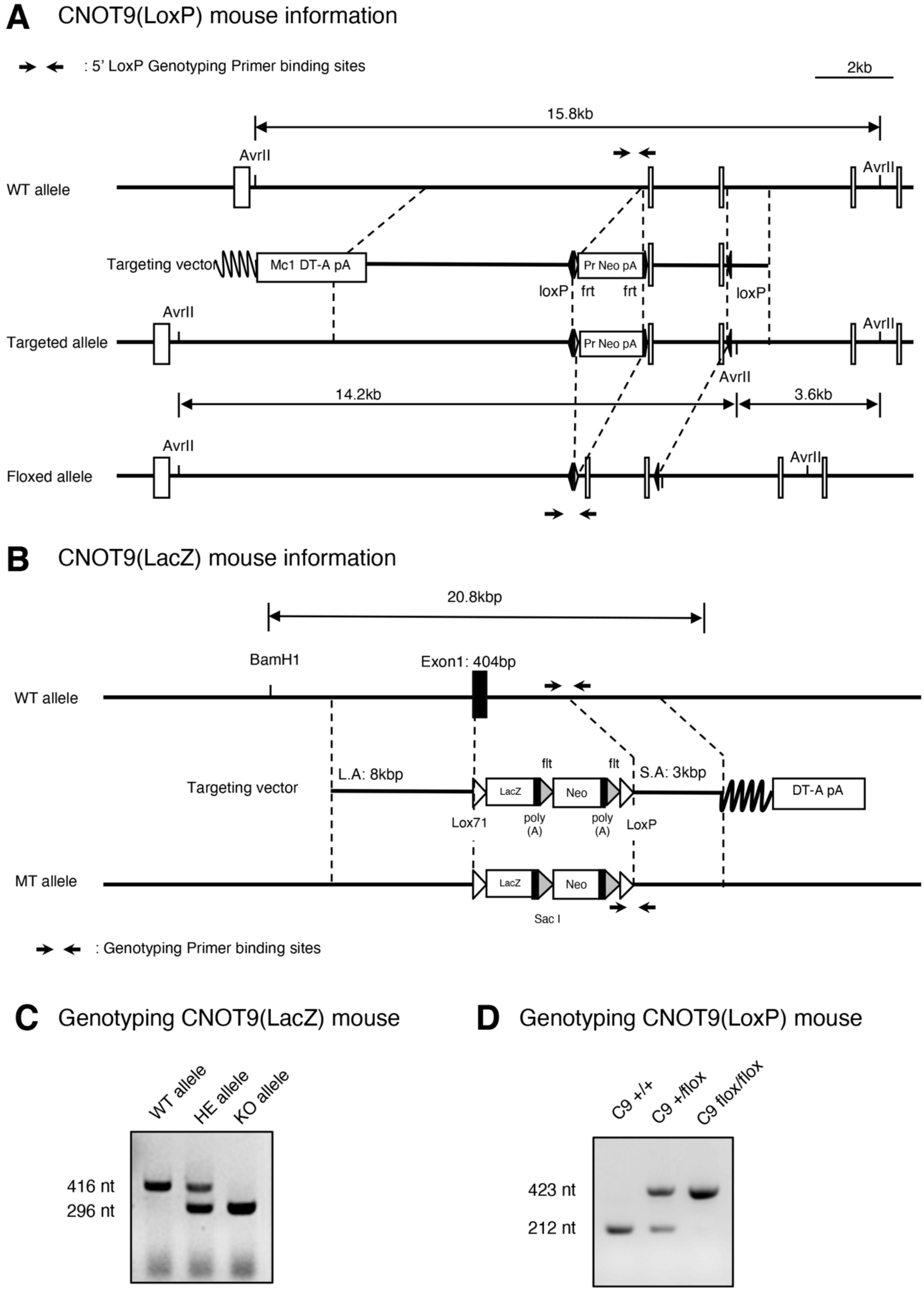


Supplementary Figure 1: Gene targeting strategy adopted for generation of **(A)** CNOT9 conditional mouse by targeting exon 2 and 3 of CNOT9 locus **(B)** LacZ knock-in mouse by targeting exon1 of CNOT9 locus. **(C)** Genotyping PCR bands corresponding to WT, HE, and KO alleles in CNOT9(LacZ) mouse **(D)** Genotyping PCR bands corresponding to WT, CNOT9(flox/+), and CNOT9(flox/flox) alleles in CNOT9(LoxP) mouse. CNOT9 is abbreviated as C9 in figure.

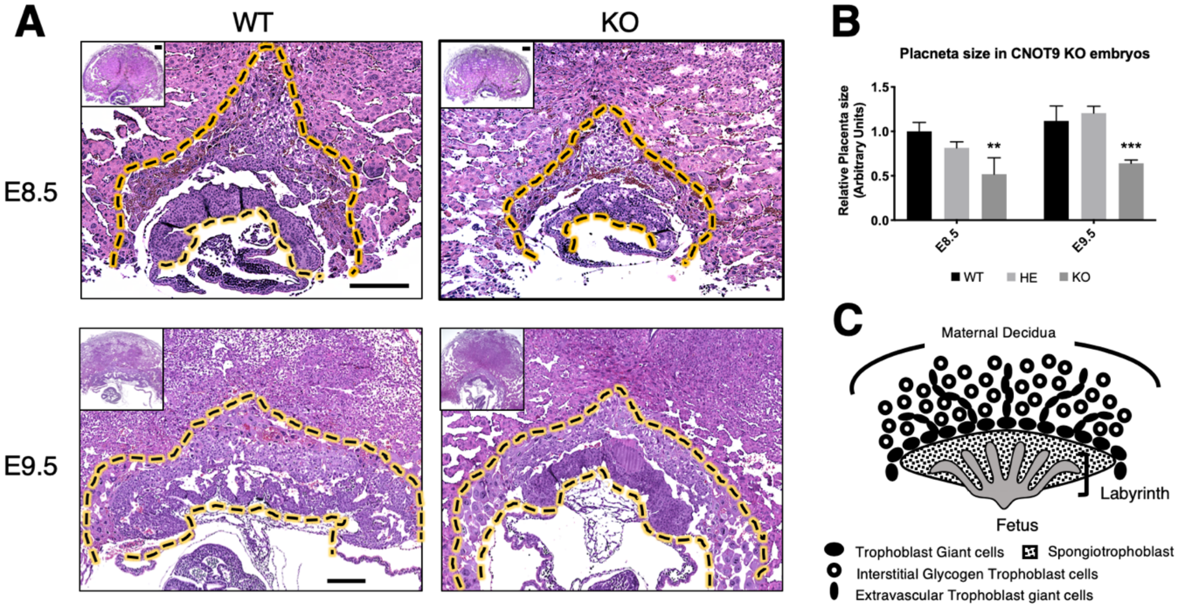


Supplementary Figure 2: **(A)** Placental morphology examined by HE staining of paraffin embedded tissue at E8.5 and E9.5 stages of embryo development. **(B)** Quantification of placenta size in CNOT9 KO embryos compared to HE and WT controls (n=3, per time point) **(C)** Schematic representation of placental lamination during E8.5-E9.5 stage. Scale: 250um

#### A Downregulated targets based on RNA-Seq data (RPKM normalized)

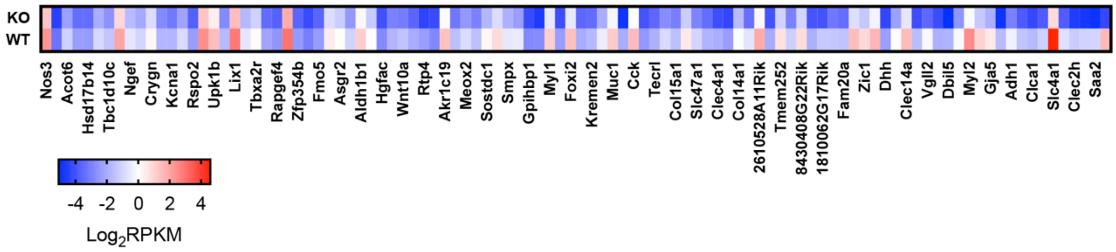


#### B Downregulated targets based validated by qRT-PCR data (GAPDH normalized)

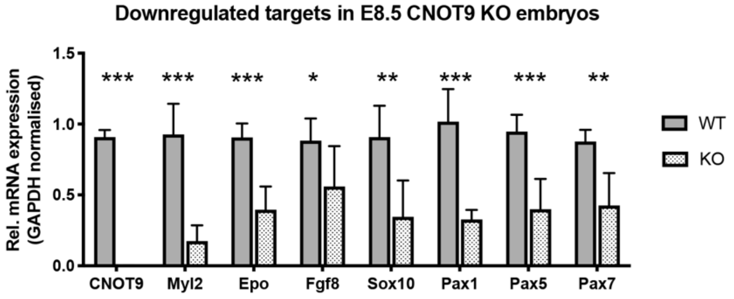


Supplementary Figure 3: **(A)** Heat-map indicating the average extent of downregulation for target genes in KO embryos compared to WT controls. (n=3) **(B)** qRT-PCR based validation of downregulated genes involved in embryonic gastrulation at E8.5 stage. Values are Mean ± SD [n=5, $p < 0.05$ (*), $p < 0.01$ (**), $p < 0,001$ (***)]

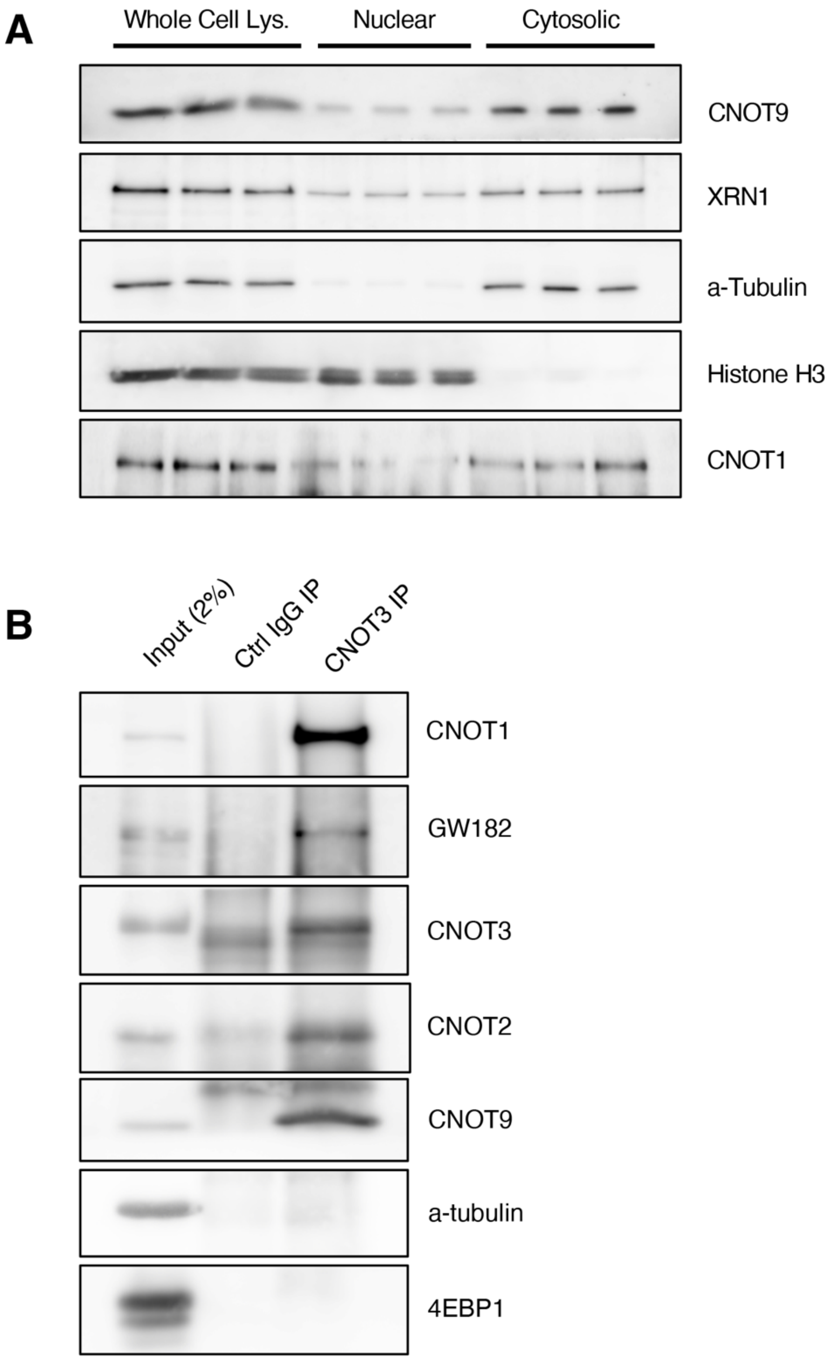


Supplementary Figure 4: (A) Nuclear-Cytoplasmic fractionation of E8.5 stage embryos to determine weightage of CNOT9 in cytosolic fraction. (B) Proteins co-immunoprecipitated using anti-CNOT3 antibody on E8.5 mouse embryo lysates.

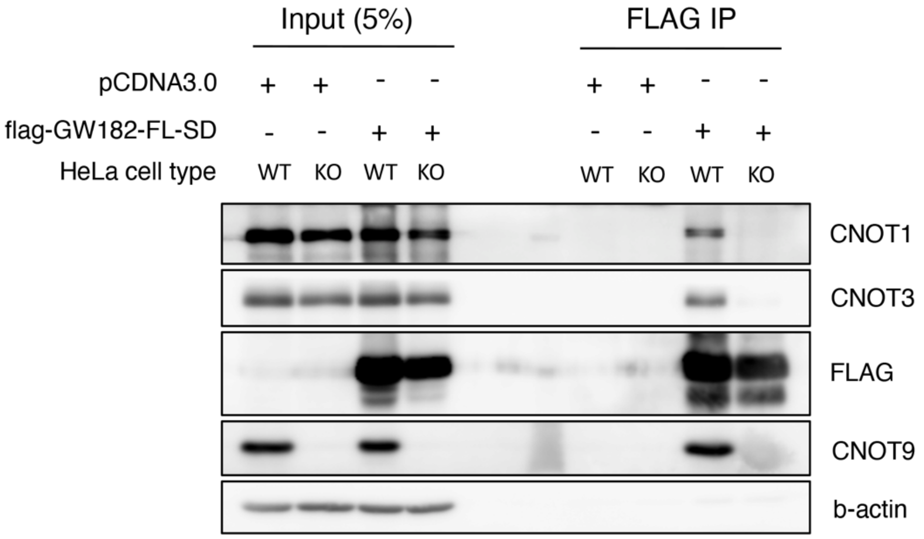


Supplementary Figure 5: Co-immunoprecipitation assay using flag-tagged silencing domain of GW182 (TNRC6C) protein as bait in WT and CNOT9 KO HeLa cells.

| Sl. No | Target | Forward Primer (5'-3') | Reverse Primer (5'-3') |
|---|---|---|---|
| 1 | *Lefty1* | actcagtatgtggccctgcta | aacctgcctgccacctct |
| 2 | *Lefty2* | gccctcatcgactctaggc | agctgctgccagaagttcac |
| 3 | *Oct4* | aagttggcgtggagactttg | tctgagttgctttccactcg |
| 4 | *Cnot9* | gtctgcgcatcatggagtc | aaccagtgtcatccaagagga |
| 5 | *Nodal* | tggtagggaagacccacaac | tgccaagcatacatctcagg |
| 6 | *Cfc-1* | tgtgttctgggcagtttctg | cctagggcaccacagtct |
| 7 | *Noto* | tgtttgcaaagcagcacaa | ttggaaccagatcctcacct |
| 8 | *Gapdh* | ctgcaccaccaactgcttag | gtcttctgggtggcagtgat |
| 9 | *Pax1* | cggacgtttatggagcaaac | tccatcttgggggagtagg |
| 10 | *Pax5* | gacgctgacagggatggt | ggggaacctccaagaatcat |
| 11 | *Pax7* | gtcccagtcttactgcccac | tgtggacaggctcacgtttt |
| 12 | *c-myb* | cctcaaagcctttaccgtacc | ctgtcttcccacaggatgc |
| 13 | *Sox10* | atgtcagatgggaacccaga | gtctttggggtggttggag |
| 14 | *Epo* | gaggcagaaaatgtcacgatg | ttccaagcatagaagttgactttg |
| 15 | *Myl2* | gtttgagcagacccagatcc | ttgtcgatgaagccgtctct |

Supplementary Table 1: List of primers used for qRT-PCR analysis
